# Neural mechanisms of speed-accuracy tradeoff of visual search: Saccade vigor, targeting errors, superior colliculus and frontal eye field

**DOI:** 10.1101/212621

**Authors:** Thomas R. Reppert, Mathieu Servant, Richard P. Heitz, Jeffrey D. Schall

## Abstract

Balancing the speed-accuracy tradeoff (SAT) is necessary for successful behavior. Using a visual search task with interleaved cues emphasizing speed or accuracy, we recently reported diverse contributions of frontal eye field (FEF) neurons instantiating salience evidence and response preparation. Here we report replication of visual search SAT performance in two macaque monkeys, new information about variation of saccade dynamics with SAT, extension of the neurophysiological investigation to describe processes in the superior colliculus, and description of the origin of search errors in this task. Saccade vigor varied idiosyncratically across SAT conditions and monkeys, but tended to decrease with response time. As observed in the FEF, speed-accuracy tradeoff was accomplished through several distinct adjustments in the superior colliculus. Visually-responsive neurons modulated baseline firing rate and the time course of salience evidence. Unlike FEF, the magnitude of visual responses in SC did not vary across SAT conditions, but the time to locate the target was longer in **Accurate** as compared to **Fast** trials. Also unlike FEF, the activity of SC movement neurons when saccades were initiated was equivalent in **Fast** and **Accurate** trials. Search errors occurred when visual salience neurons in FEF and SC treated distractors as targets, even in the **Accurate** condition. Saccade-related neural activity in SC but less FEF varied with saccade peak velocity. These results extend our understanding of the cortical and subcortical contributions to SAT.

**Significance statement:** Neurophysiological mechanisms of speed-accuracy tradeoff (SAT) have only recently been investigated. This paper reports the first replication of SAT performance in nonhuman primates, the first report of variation of saccade dynamics with SAT, the first description of superior colliculus contributions to SAT, and the first description of the origin of errors during SAT. These results inform and constrain new models of distributed decision-making.

## Introduction

The speed-accuracy tradeoff (SAT) is a fundamental behavioral phenomenon (Heitz, 2014). Computational decision models explain SAT in terms of a stochastic accumulation of noisy sensory evidence from a baseline level over time; responses are produced when the accumulated evidence for one choice reaches a threshold. Slower, more accurate responses are achieved by elevating the threshold; faster, less accurate responses are produced by lowering the threshold. Many laboratories have provided evidence linking the stochastic accumulation process with the activity of specific neurons in the frontal eye field (FEF) (Boucher et al., 2007; Ding and Gold, 2012; Hanes and Schall, 1996; Kim and Shadlen, 1999; Purcell et al., 2010, 2012a; Woodman et al., 2008), lateral intraparietal area (LIP) (Roitman and Shadlen, 2002; Wong et al., 2007); but see (Latimer et al., 2015; Yates et al., 2017), motor and premotor cortex (Thura et al., 2012), superior colliculus (SC) (Ratcliff et al., 2003, 2007) and basal ganglia (Ding and Gold, 2010). However, neurophysiological studies investigating SAT have only recently appeared (Hanks et al., 2014; Heitz and Schall, 2012; Thura and Cisek, 2016, 2017), and all have focused on forebrain structures. We now replicate major findings in two more monkeys and show that subcortical processes in the superior colliculus also contribute to SAT for saccades during visual search. We also show how search errors arise in this task and report new findings about idiosyncratic and systematic variation of saccade dynamics across SAT conditions.

## Methods

All procedures were approved by the Vanderbilt Institutional Animal Care and Use Committee in accordance with the United States Department of Agriculture and Public Health Service Policy on Humane Care and Use of Laboratory Animals.

### Task

Four bonnet macaques (*M. radiata*), identified as *Q, S, Da* and *Eu*, performed a form (**T/L**) visual search task for a target item presented amongst seven distractor items (Figure 1). Trials began when monkeys fixated a central cue for approximately 1 sec. Each monkey was extensively trained to associate the color of the fixation cue (red or green) with a task condition (**Accurate** or **Fast**). After fixation, an iso-eccentric array of **T/L** shapes appeared, of which one was the target item for that day. Distractor items were drawn randomly from the non-target set and oriented randomly in the cardinal positions.

**Figure 1.**
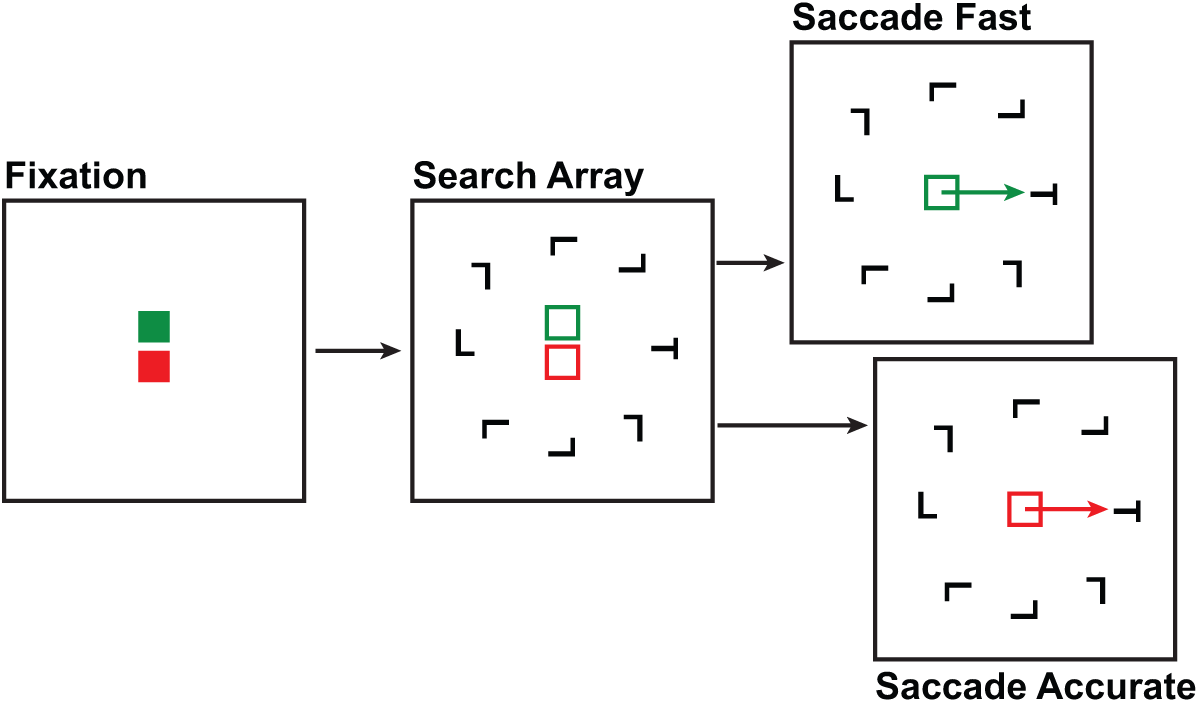
Visual search paradigm with speed-accuracy tradeoff. Trials began with a fixation cue signifying whether the trial was to be **Fast** (green) or **Accurate** (red). After a fixation period of ~ 1 sec, an iso-eccentric array of 8 T/L shapes appeared, of which one was the target for that session. Monkeys searched for the target item (rotated T or L) presented with 7 distractors (rotated L or T). In the **Fast** and **Accurate** conditions, correct responses were only rewarded when executed before and after an unsignaled deadline, respectively. In some sessions, distractors were of homogeneous orientation; in other sessions they were randomly rotated. Fixation point, search shapes, and viewing screen are not drawn to scale.

Sessions were comprised of mini-blocks of 10-20 trials of a single SAT condition. In the **Accurate** condition, saccades to the target item were rewarded if RT exceeded an unsignaled deadline. The deadline was adjusted to optimize the yield of useful data against sustained performance. In this data set the deadline for successful trials in the **Accurate** condition was set to, respectively, 500 ms (*Q*), 427 ± 5 ms (*S*), 435 ± 7 ms (*Da*), and 445 ± 6 ms (*Eu*) (mean ± SEM across sessions). In the **Accurate** condition, responses with incorrect timing and/or direction were followed by a 4-sec time-out. In the **Fast** condition, saccades to the target item were rewarded only if the saccade to the target occurred before an unsignaled deadline. In this data set the deadline for successful trials in the **Fast** condition was, respectively, 370 ± 10 ms (*Q*), 386 ± 7 ms (*S*), 365 ± 14 ms (*Da*), and 429 ± 21 ms (*Eu*) (mean ± SEM). Saccades executed after the deadline in the **Fast** condition were followed by a 4-sec time-out. However, inaccurate saccades made before the deadline in the **Fast** condition (i.e. with correct timing) had no time-out. The lack of timeout following mis-directed saccades was used to incentivize quick responses.

### Neural data acquisition

Details of the methods have been reported previously (Heitz & Schall, 2012). Briefly, neural spikes were sampled in SC and FEF using tungsten microelectrodes (2-4 MΩ, FHC, Inc.). Location was verified by evoking eye movements with < 50 µA electrical micro-stimulation. The number of electrodes lowered during a given session ranged from 1 to 9. Single unit waveforms were digitized and sorted offline (Offline Sorter; Plexon Inc., Dallas, TX).

### Neural data analysis

Spike trains were convolved with a kernel that resembled a post-synaptic potential to create a spike density function (SDF) (τ_growth_ = 1 ms, τ_decay_ = 20 ms, Thompson et al., 1996). For visually-responsive activity, SDFs were normalized to the peak average activity during the 1-sec time interval post-stimulus appearance. For saccade-related activity, SDFs were normalized to the peak average activity in the time interval 100 ms pre-to 100 ms post-saccade initiation. Normalization factors were computed across all conditions and behavioral outcomes (i.e., over all SAT conditions, all RT, correct and errant responses, etc.) in a particular session.

Neurons were categorized into 3 major types based on gross function: visual, visuo-movement, and movement. Though classification operates along a continuum, many observations demonstrate that these populations are functionally distinct (Ray et al., 2009; Cohen et al., 2009b; Gregoriou et al., 2012). Visual neurons increase discharge rates significantly immediately following search array presentation but have little or no saccade-related modulation. Movement neurons increase discharge rate significantly before saccade initiation but have little or no visual response. Visuo-movement neurons exhibit both periods of modulation. To classify neurons, we analyzed single-unit activity from a memory-guided saccade task. The primary function of this task was to dissociate visual activity related to stimulus appearance from movement activity related to saccade execution. We compared activity in the interval after stimulus presentation to activity preceding presentation. All neurons with higher activity in the post-presentation interval were classified as visual neurons. Similarly, we compared activity in the interval immediately before saccade initiation to activity 500-400 ms before the saccade. All neurons with higher activity immediately prior to saccade initiation were classified as movement neurons.

### Behavioral analyses

All gaze data were collected with video-oculography (Eyelink, SR Research) at a sampling rate of 1000 Hz. The data were filtered with a 3^rd^-order Butterworth low-pass filter with a cutoff frequency of 80 Hz. The data were then differentiated to velocity traces, and a cutoff velocity of 20˚/sec was used to identify all saccades. Any movements that did not meet the following criteria were removed from all analyses: (1) duration between 10 and 80 ms, (2) peak velocity between 200 and 1000˚/sec, and (3) displacement greater than 2.5˚.

### Saccade vigor

To characterize saccade vigor, we quantified the relationship between displacement and peak velocity of all task-relevant saccades. This relationship is typically linear for saccades of displacement up to ~20˚ (Bahill et al., 1975; Collewijn et al., 1988). The majority of saccades recorded from all monkeys shifted gaze less than 9˚. Therefore, we fitted the relationship between displacement and peak velocity to a linear function constrained to pass through the origin: *g(x) = α • x*. Given the average main sequence relationship, we then computed the vigor of saccade *j* with displacement *x* as the ratio between the measured velocity and the expected velocity: 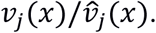 Ratios > 1.0 measure saccades with greater vigor than the saccade population average. We used this measure of vigor to quantify changes in saccade dynamics across SAT conditions and response times.

### Statistical analyses

All t-tests and Mann-Whitney U-tests are two-sided, unless otherwise stated. Corresponding test statistics and p-values are provided. To test the interaction between RT and error rate, and RT and saccade vigor, we use repeated-measures ANOVA. To determine when neurons responded differently to two SAT conditions, or when the target as compared to distractors appeared in the RF, we implement a ms-by-ms one-sided Mann-Whitney U-test, against the null hypothesis that Target-in-RF activity is not different from Distractor-in-RF activity. Target selection time (TST) is the first successive 30 ms with significant difference at the *p* < 0.01 level. We report Bayes Factor with default *r* scale of 0.707 to supplement null findings regarding visually-responsive and saccade-related activity.

## Results

### Response time and accuracy

An analysis of the performance of *Q* and *S* was previously published (Heitz and Schall, 2012). Performance measures of *Da* and *Eu* were collected during 16 sessions (*Da* – 9 sessions; *Eu* – 7). Monkeys *Da* and *Eu* adjusted response time (RT) in accordance with task condition in each session. Average RT across sessions during **Fast** condition was 280 ± 9 ms (*Da*) and 354 ± 13 ms (*Eu*) (mean ± SEM, Fig. 2A). Average RT during **Accurate** condition was 498 ± 9 ms (*Da*) and 510 ± 17 (*Eu*). Monkeys *Da* and *Eu* adjusted RT immediately upon receiving a new SAT condition cue (Fig. 2B), replicating the performance of monkeys *Q* and *S*.

**Figure 2.**
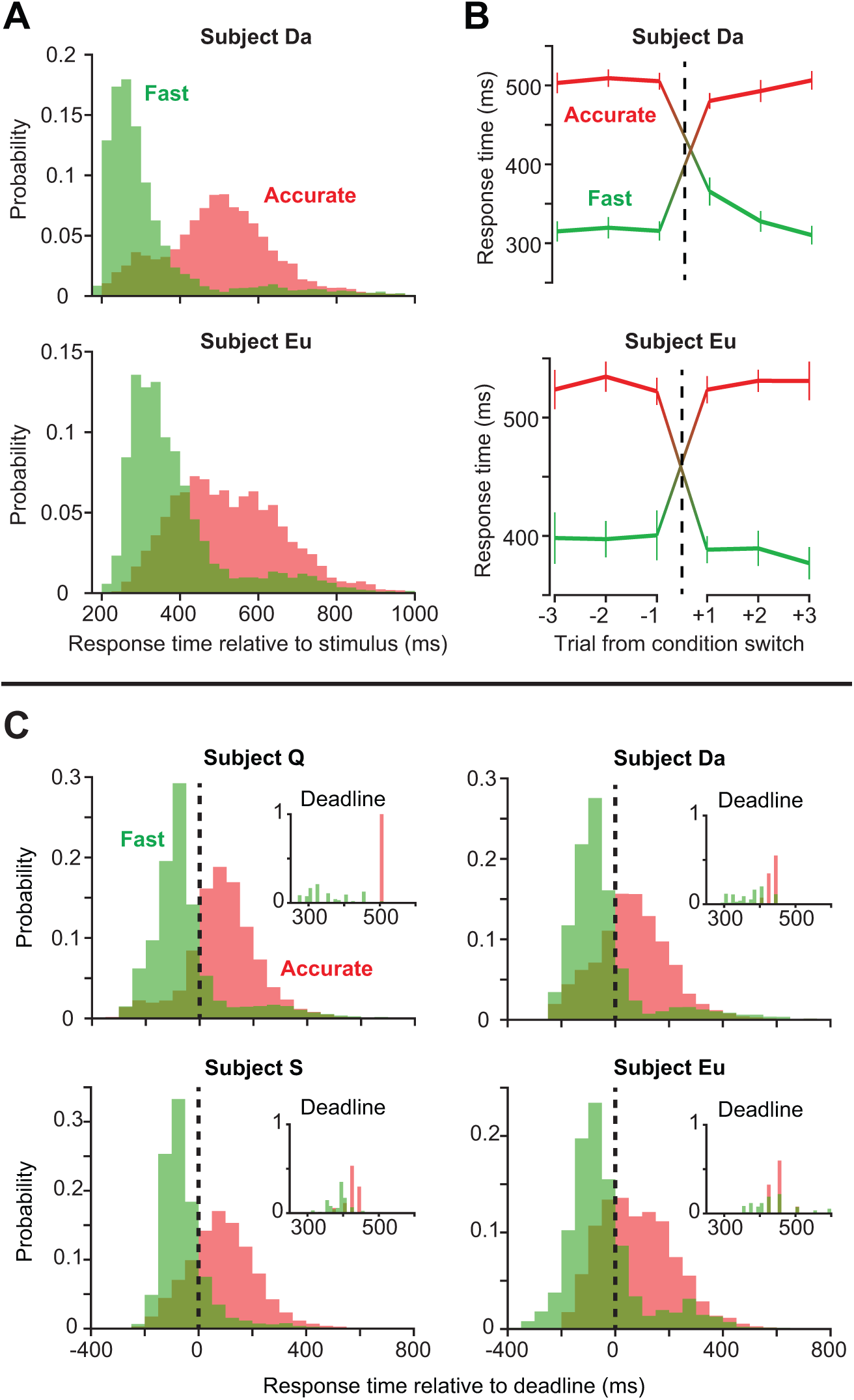
Response time adjustments during speed-accuracy tradeoff. (**A)** Probability distributions of RT on **Fast** (green) and **Accurate** (red) trials for monkeys *Da* and *Eu*. Distributions include trials pooled across all recording sessions for each monkey. (**B**) Change in RT on 3 trials immediately preceding and following a switch in task condition. Both monkeys altered RT on the trial immediately following a change in fixation cue color, signaling condition switch. Error bars are SEM. (**C**) Probability distributions of RT relative to response deadline for all 4 monkeys. Insets show probability distributions of response deadline. All responses that occurred after the deadline in the **Fast** condition or before the deadline in the **Accurate** condition were considered timing errors.

The **Accurate** and **Fast** conditions employed response deadlines similar to some human studies (Heitz and Engle, 2007; Rinkenauer et al., 2004). These deadlines were adjusted so that ~70-80% of saccades would be executed with the correct timing. Figure 2C presents RT distributions relative to the deadlines for all 4 monkeys. The prior report of performance data from *Q* and *S* did not include this measure, but these distributions support future computational modeling.

In addition to timing errors, we also analyzed saccade endpoint errors (i.e. gaze shifts to a non-target item). Average error rates across sessions in the **Fast** condition were 26% (*Da*, Fig. 3A) and 33% (*Eu*), whereas those in the **Accurate** condition were 11% (*Da*) and 14% (*Eu*). These error rates were comparable to those observed previously with monkeys *Q* and *S*. Late responses were observed in 22% (*Q*), 17% (*S*), 20% (*Da*), and 26% (*Eu*) of trials in the **Fast** condition. Premature responses were observed in 22% (*Q*), 23% (*S*), 31% (*Da*), and 32% (*Eu*) of trials in the **Accurate** condition.

**Figure 3.**
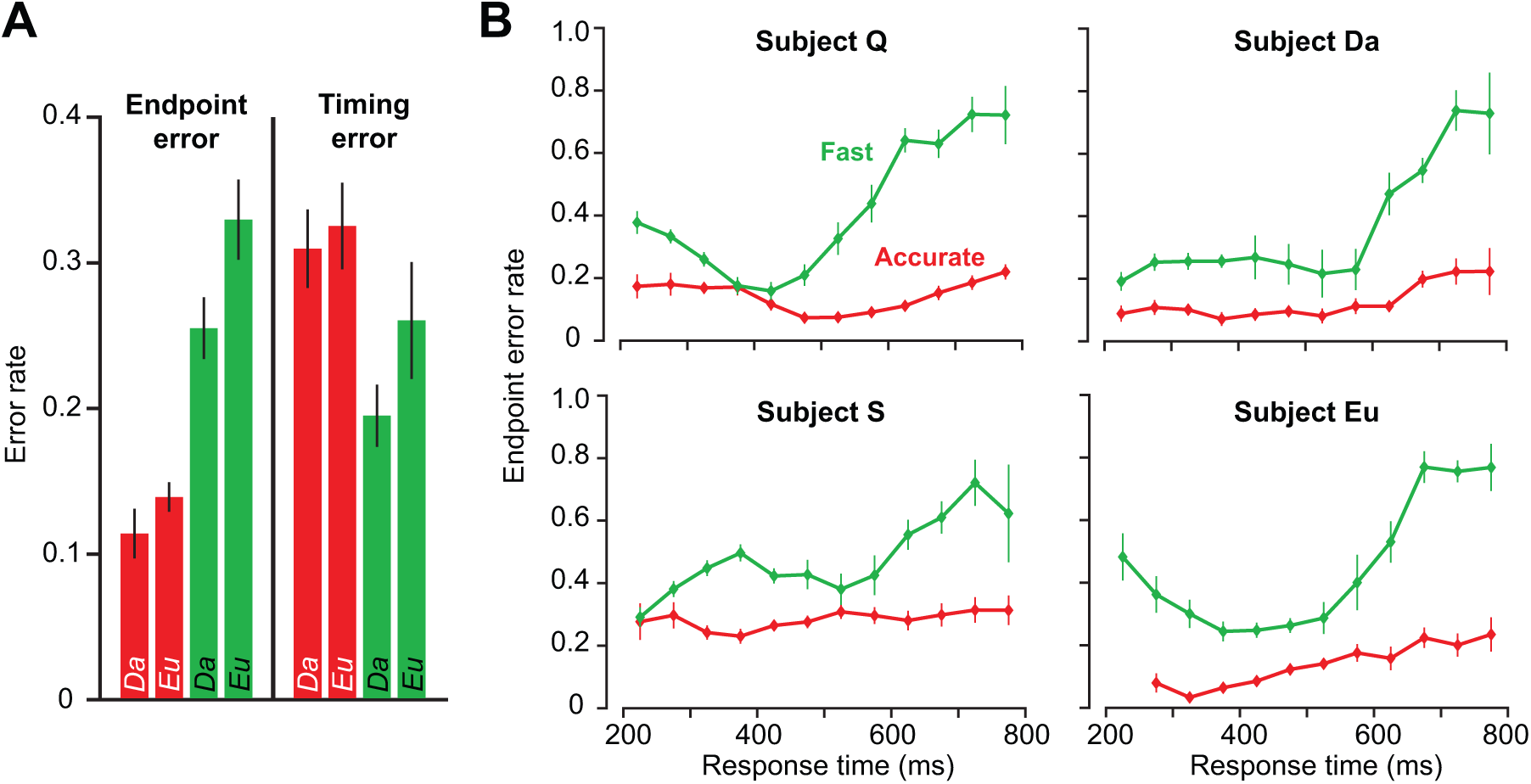
Saccade endpoint and timing errors. (**A**) Average error rates in **Fast** and **Accurate** conditions for monkeys *Da* and *Eu*. In the **Fast** condition, the monkeys committed more endpoint errors. Conversely, in the **Accurate** condition, the monkeys committed more timing errors. Monkey *Da* generally committed fewer errors than monkey *Eu*. (**B**) Change in endpoint error rate with RT. We binned endpoint error rate by condition and RT, and then averaged binned endpoint error rate across all sessions for each monkey. Error rate increased by ~ 40-50 % during the **Fast** condition. Error bars are SEM.

As the monkeys adjusted RT to accommodate timing criteria in **Fast** and **Accurate** conditions, saccade error rates changed accordingly. We asked whether endpoint error rates changed with RT in either condition. For each monkey, we binned trials by RT, and calculated the probability that an endpoint error occurred for each RT bin (Fig. 3B). We then averaged this binned endpoint error rate across all sessions for each monkey. We found that endpoint error rate increased during the **Fast** condition (repeated-measures ANOVA, *Q*: F_(5,105)_ = 31.0, p = 6×10^−19^; *S*: F_(5,75)_ = 9.7, p = 3×10^−7^; *Da*: F_(5,40)_ = 23.3, p = 7×10^−11^; *Eu*: F_(5,30)_ = 21.2, p = 5×10^−9^). We also found a significant change in endpoint error rate during the **Accurate** condition (*Q*: F_(5,105)_ = 8.0, p = 2×10^−6^; *S*: F_(5,80)_ = 2.5, p 0.04; *Da*: F_(5,40)_ = 7.3, p = 6×10^−5^; *Eu*: F_(5,30)_ = 15.7, p = 1×10^−7^).

However, the average increase in error rate during **Accurate** condition (~10%) was much less than the average increase during the **Fast** condition (~50%).

### Saccade vigor

Heitz & Schall (2012) motivated an interpretation of the pattern of modulation in FEF across SAT conditions by the observation that saccade velocities seemed invariant across conditions, and Heitz & Schall (2013) showed such data from a single session. Here, we examine this issue with the more sensitive measure of saccade vigor (Choi et al., 2014; Reppert et al., 2015). We fitted Eq. (1) to the saccade main sequence for each monkey across all task-relevant saccades, producing a monkey-specific mean slope α (74.03 ± 0.09 (*Q*); 70.28 ± 0.12 (*S*); 71.54 ± 0.12 (*Da*); and 68.71 ± 0.16 (*Eu*), mean ± 95% CI) with high goodness of fit for each monkey: R^2^ = 0.75 (*Q*); 0.67 (*S*); 0.81 (*Da*); and 0.59 (*Eu*). This one-parameter model accounted for, on average, 71% of the variance in the monkeys’ saccade peak velocities.

We first asked whether the velocity profile of saccades changed with RT. For each session, we split all task-relevant saccades by condition and RT quartile, and plotted the average profile for the 1^st^ and 4^th^ quartiles (Fig. 4A, **Fast** condition). We focused this analysis on saccades with displacement between 5.5 and 6.5 deg, given that most task-relevant saccades were of this magnitude. We found that, during the **Fast** condition, peak velocity of saccades decreased with RT for all 4 monkeys (Fig. 4B, t_(18)_=13.2, p=1×10^−10^ (*Q*), t_(16)_=7.6, p=1×10^−6^ (*S*), t_(5)_=2.6, p=0.049 (*Da*), t_(6)_=2.7, p=0.038 (*Eu*), paired t-test). We observed no equivalent effect of RT on peak velocity in the **Accurate** condition (p=0.74 (*Q*), p=0.14 (*S*), p=0.19 (*Da*), p=0.45 (*Eu*)).

**Figure 4.**
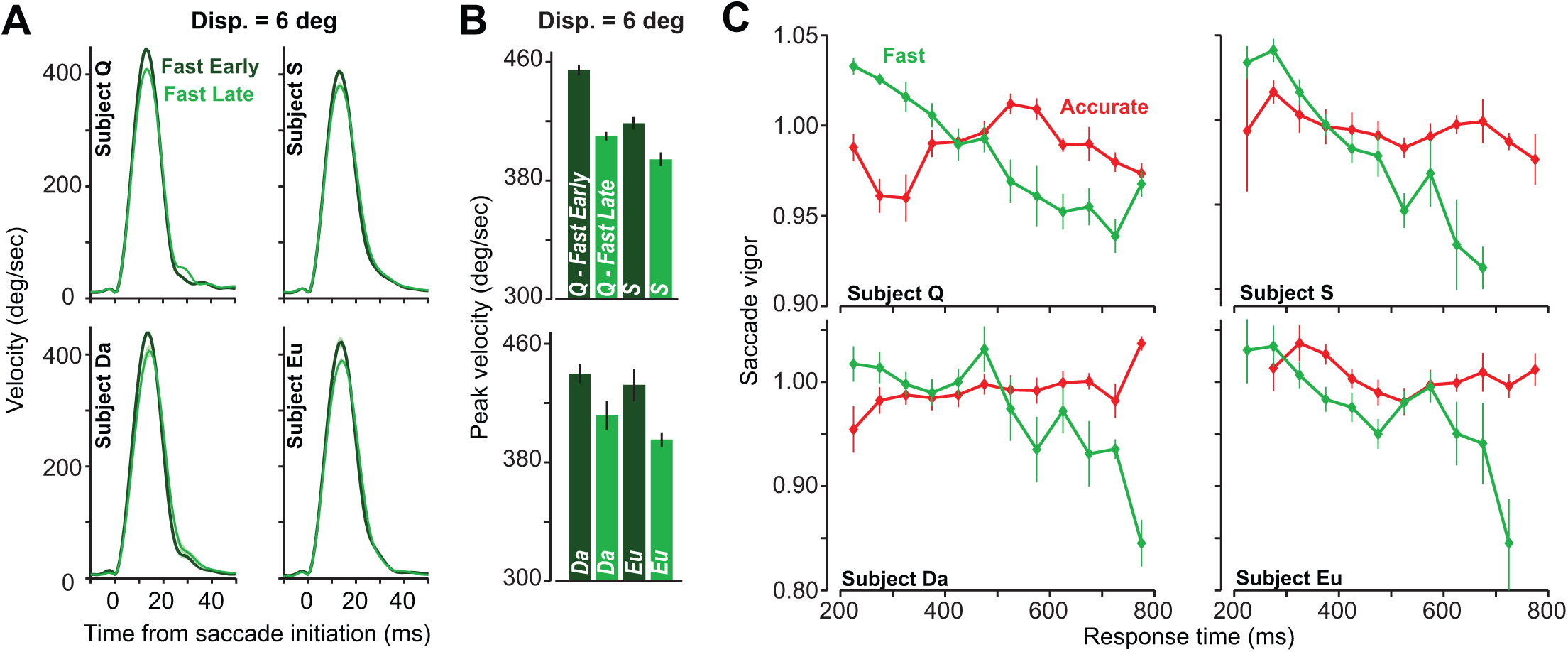
Saccade vigor. (**A**) Average velocity profiles for early (dark green) and delayed (light green) responses in the **Fast** condition. For each recording session, we computed RT quartiles, and average velocity profiles for the responses comprising the 1^st^ and 4^th^ quartiles of the RT distribution. We then averaged these profiles across recording sessions. Profiles are shown for responses with displacement between 5.5 and 6.5 deg. (**B**) Average peak velocity for early and late responses in the **Fast** condition. As in (A), saccades belong to the 1^st^ and 4^th^ RT quartiles of the recording session. Peak velocity decreased with RT during the **Fast** condition for all 4 monkeys. (**C**) Change in saccade vigor with RT. We binned trials by RT, and calculated the average vigor of saccades for each bin. We then averaged binned vigor across all sessions. Vigor of saccades decreased with RT in the **Fast** condition, and varied idiosyncratically in the **Accurate** condition.

Did this decrease in saccade peak velocity exist for saccades of all displacements? To answer this question, we defined vigor of saccade *j* as the ratio of peak velocity to expected peak velocity: 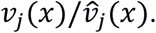 This method afforded us a comparison of velocities across saccade displacements. We then asked whether saccade vigor changed with RT in the **Fast** and **Accurate** conditions (Fig. 4C). We found a significant decrease of saccade vigor with prolonged RT in the **Fast** condition (repeated-measures ANOVA, *Q*: F_(5,95)_ = 16.4, p = 1×10^−11^; *S*: F_(5,55)_ = 6.2, p = 1×10^−4^; *Da*: F_(5,25)_ = 3.5, p = 0.02; *Eu*: F_(5,25)_ = 6.7, p = 4×10^−4^). Vigor decreased with RT in the **Fast** condition on the order of ~10% (*Q*), 10% (*S*), 15% (*Da*), and 15% (*Eu*). On average, saccade vigor decreased ~12.5% from early to late RT during the **Fast** condition. We found no such effect of RT on vigor for the **Accurate** condition.

### Introduction of neural discharge data

The monkeys mastered the fixation-cued SAT search task well enough to alter RT immediately when the SAT cue changed (Fig. 2B). Similarly, adjustments in behavior were accompanied by changes in firing rate so pronounced that they were visible in the spike rasters (Fig. 5A, monkey *Q*). Importantly, these changes in firing rate were visible even during the fixation period in some FEF neurons. As the monkeys adjusted behavior to satisfy the **Fast** or **Accurate** instruction, neurons in FEF and SC demonstrated markedly different profiles of activity. The two neurons illustrated were recorded simultaneously in FEF. The parallel modulation of both neurons indicates that SAT is accomplished with a global state change influencing the networks in FEF and SC.

**Figure 5.**
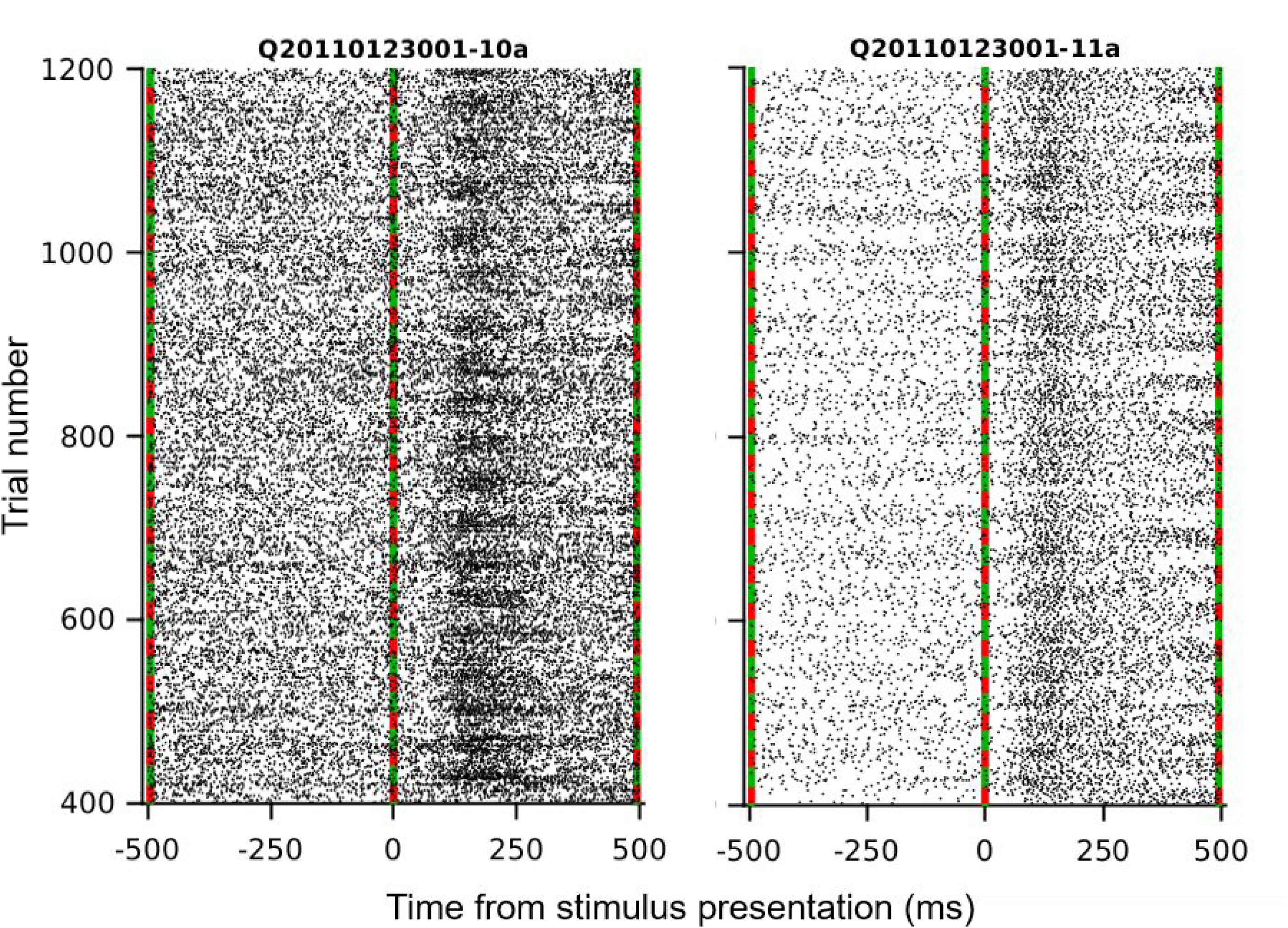
Speed-accuracy modulation of discharge rates in FEF. Rasters plot spike times relative to array presentation for two neurons recorded simultaneously in FEF on separate electrodes during the same session. A subset of 800 trials is plotted. Trials alternated between ~20 trial blocks of **Fast** (green) and **Accurate** (red) conditions. Pronounced effects of SAT cuing were visible both before and after array presentation.

As observed in FEF, SAT of visual search is accomplished by multiple adjustments in the activity of distinct types of neurons in SC. We will first describe the adjustments in baseline activity (i.e. activity prior to stimulus appearance) of samples of 22 visually-responsive FEF neurons (*Da*), and 10 visually-responsive SC neurons (*Da* n=6; *Eu* n=4). These neurons increased firing rate when salient items appeared in their receptive field (RF). We will assess response magnitude and target selection time (TST) in both sets of neurons. Visual search decisions are guided by this representation of stimulus salience in FEF (Purcell et al., 2010, 2012a; Sato and Schall, 2003; Sato et al., 2001; Thompson et al., 1996), posterior parietal cortex (Balan et al., 2008; Buschman and Miller, 2007; Constantinidis and Steinmetz, 2005; Gottlieb et al., 1998; Ipata et al., 2006; Ogawa and Komatsu, 2009; Thomas and Paré, 2007), substantia nigra pars reticulate (Basso and Wurtz, 2002), ocular motor thalamic nuclei (Wyder et al., 2004), and SC (Kim and Basso, 2008; McPeek and Keller, 2002; Shen and Paré, 2007; White and Munoz, 2011). Data from visual and visuo-movement neurons were combined for analyses of visual response activity. On error trials, the visually-responsive neurons exhibited increased firing rate when a distractor was placed in the neuron’s RF. We will also report for the first time the origin of search errors in both **Fast** and **Accurate** conditions. This analysis will include data that was previously reported (Heitz and Schall, 2012) (*Q* n=83, *S* n=90; visually-responsive neurons).

We will extend our assessment of SAT-related neural activity with a description of saccade-related buildup activity in SC neurons (*Da* n = 2, *Eu* n = 4) and FEF neurons (*Da* n = 18) that have been identified with stochastic accumulation of the salience evidence provided by visual neurons (Ratcliff et al., 2003, 2007). Pure movement neurons – those with no visual response – are encountered less commonly than neurons with visual responses. Here, we only recorded from 1 pure movement-related neuron in SC, and 2 pure movement neurons in FEF. We will conclude with an analysis of the relationship of the activity of FEF and SC movement neurons with saccade velocity.

### Adjustments in baseline activity

Heitz and Schall (2012) previously reported significant modulation of baseline activity in 54% of FEF neurons with visually-responsive activity. The majority of these neurons exhibited elevated activity in the **Fast** condition. We tested these findings in a set of 22 visually-responsive FEF neurons (*Da*, Fig. 6A). Thirteen out of 22 neurons (59%) exhibited significant modulation of activity during the baseline period before array stimulus appearance (Mann-Whitney U-test, p<0.05). Nine of these 13 neurons (69%) exhibited elevated activity during baseline in the **Fast** condition. Despite this effect for single neurons, we observed no significant effect of task condition on baseline activity across the population of 22 visually-responsive FEF neurons (Fig. 6C, paired t-test, t_(22)_ = 0.70, p = 0.49, BF = 3.7). What was the timecourse of change in baseline activity for those neurons that did exhibit a condition-cued shift? We assessed trial-to-trial changes in baseline activity around the time of switch in condition. Baseline activity of each neuron was normalized to average firing rate during the 750-ms interval prior to array presentation. Baseline activity in FEF responded to a change in task condition within a single trial (Fig. 6E).

**Figure 6.**
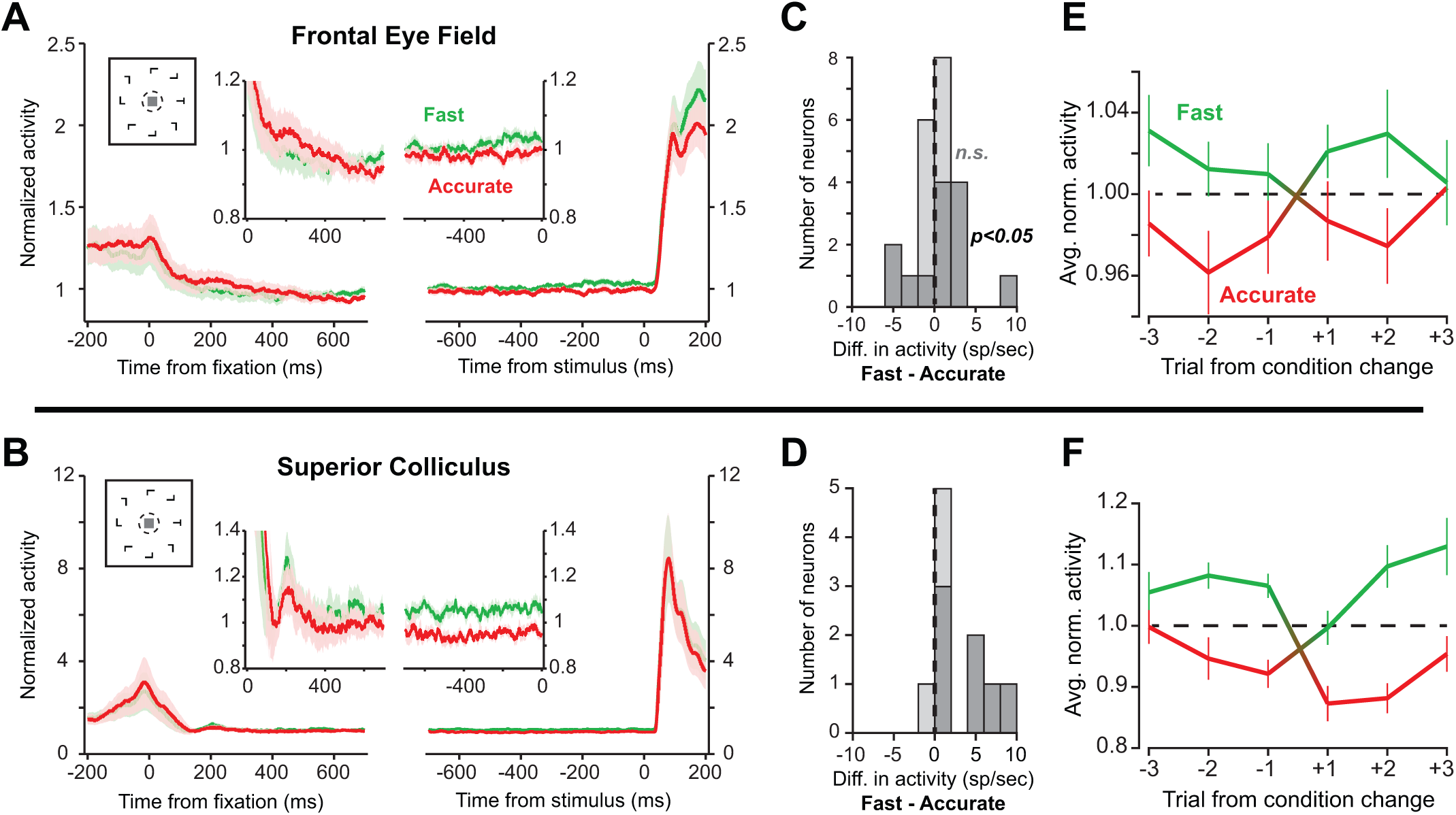
Adjustment of baseline activity during speed-accuracy tradeoff. (**A**) Average baseline activity of 22 visually-responsive FEF neurons. Plots on left and right are aligned on fixation of the central cue stimulus, and appearance of array stimulus ~ 1 sec later. Diagrammatic inset shows fixation of central cue upon appearance of array stimulus. Function-based inset shows baseline activity in both conditions. Baseline activity was elevated during the **Fast** condition. Error regions are SEM. (**B**) Average baseline activity for 10 visually-responsive SC neurons. Baseline activity was elevated during the **Fast** condition. Conventions as in (A). (**C**) Histogram of difference in baseline firing rate of FEF neurons during **Fast** and **Accurate** conditions. Dark and light bars mark neurons with and without a significant change in baseline activity (Mann-Whitney U-test, p < 0.05). (**D**) Histogram of difference in average baseline firing rates of SC neurons. Conventions as in (C). (**E**) Trial-to-trial change in baseline activity of visually-responsive FEF neurons at a fixation-cued change in condition. Activity of each neuron was normalized to average baseline activity across all trials, both **Fast** and **Accurate**. Error bars are SEM. (**F**) Trial-to-trial change in baseline activity of visually-responsive SC neurons at a fixation-cued change in condition. Conventions as in (E).

Were the effects of SAT-cued speed constraints a global phenomenon? We next assessed baseline activity in 10 visually-responsive SC cells (*Da* n = 6; *Eu* n = 4). We found that SAT cue induced a shift in baseline firing rates following fixation and preceding array presentation in visually-responsive SC neurons (Fig. 6B). Across the population of 10 SC neurons, we found 7 neurons (3 pure visual, 4 visuo-movement) with significantly elevated baseline activity in the **Fast** condition (p<0.05, Mann-Whitney U-test). Across the population, baseline activity was significantly elevated during the **Fast** condition (Fig. 6D, paired t-test, t_(9)_=2.67, p = 0.03). As for FEF, we assessed the of trial-to-trial changes in baseline activity in SC. Normalized baseline activity responded to a change in task condition within a single trial (Fig. 6F). Neurons with and without baseline modulation were recorded within single sessions and even single electrode penetrations. Thus, SAT is accomplished in part through an immediate adjustment of cognitive state before stimuli are presented.

In summary, we observed elevated baseline activity in the **Fast** condition in SC (monkeys *Da* and *Eu*). However, while some visually-response FEF neurons did show elevated activity in **Fast** condition, this result was not consistent across the population. Cued changes in baseline activity occurred within a single trial after a switch from **Fast** to **Accurate** or **Accurate** to **Fast** condition.

### Evidence representation adjustments

Visually-responsive neurons increase firing rate when salient items appear in their receptive field (RF). All 10 visually-responsive SC neurons distinguished target items from distractors by firing at a higher rate after an initial nonselective period (Fig. 7A; McPeek and Keller, 2002; Shen and Paré, 2007; White et al., 2017). We asked whether response speed constraints caused a shift in TST in **Fast** and **Accurate** conditions. For each neuron, TST was estimated using a sliding-window ms-by-ms Mann-Whitney U-test (first 30-ms period, p < 0.01) on single-trial SDFs. The average TST for SC neurons in the two conditions was 173.5 ms (**Fast**) and 226.2 ms (**Accurate**). We found a significant effect of speed constraints on TST, with earlier TST in the **Fast** condition (Fig. 7C, paired t-test, t_(9)_ = 2.63, p = 0.03).

**Figure 7.**
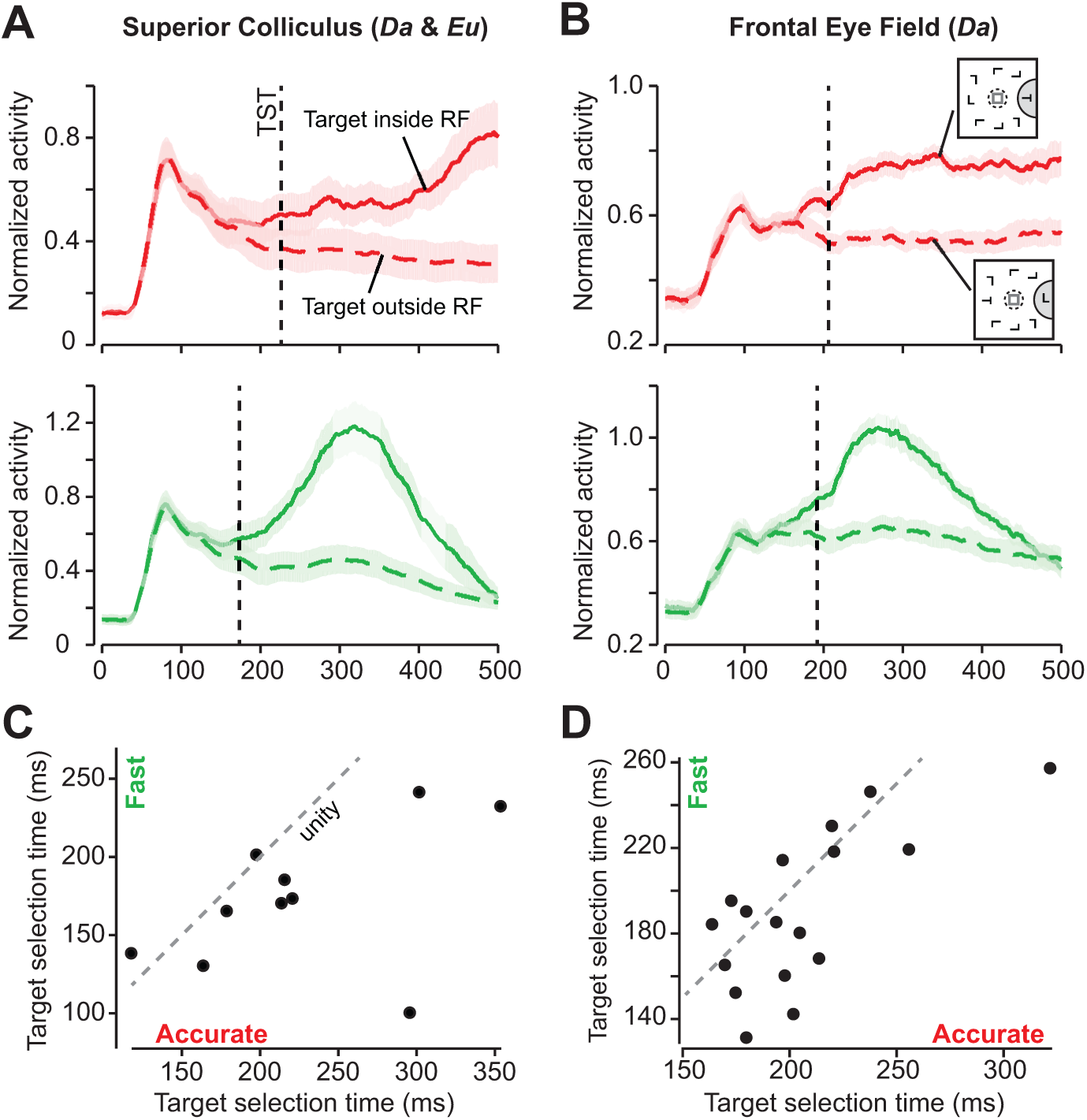
Effects of speed-accuracy tradeoff on salience evidence. (**A**) Average normalized SDF of visually-responsive SC neurons (*Da* n=6, *Eu* n=4) during the 500-ms interval following presentation of the array stimulus. Density functions for **Accurate** (*top*) and **Fast** (*bottom*) conditions are shown separately. Each neuron’s activity was normalized to maximum average firing rate during the 1-sec interval post-array appearance. Only trials with correctly placed and timed responses were included. Dashed lines show average TST (174 ms and 226 ms on **Fast** and **Accurate** trials, respectively). Error regions are SEM. (**B**) Average normalized SDF of all visually responsive FEF neurons that selected the target amongst distractors (*Da* n=17). Diagrammatic inset shows schematic of the search array on trials when the target was placed inside (*top*) and outside (*bottom*) of the neuron’s RF. Average TST was 190 ms and 206 ms on **Fast** and **Accurate** trials, respectively. (**C**) Neuron-wise relationship between TST in the **Fast** and **Accurate** conditions for SC. Each dot represents a single neuron. Dashed line is unity. Most SC neurons had earlier TST in the **Fast** condition (**D**) Neuron-wise relationship between TST in **Fast** and **Accurate** conditions for FEF. Most FEF neurons had earlier TST in the **Fast** condition.

Heitz and Schall (2012) previously found that FEF neurons selected the target amongst distractors earlier in the **Fast** condition. We tested this finding in our sample of visually-responsive FEF neurons. Seventeen of these 22 neurons (77%) distinguished target items from distractors (Fig. 7B). The average TST of visually-responsive FEF neurons in the two conditions was 190.4 ms (**Fast**) and 206.4 ms (**Accurate**). Across our sample, TST was significantly earlier in the **Fast** condition (Fig. 7D, t_(16)_ = 2.28, p = 0.04), replicating the prior findings (2012).

We also asked whether SAT task constraints affected the magnitude of the visual response to array stimulus in SC or FEF. For each task condition, we computed the maximum value of the visual response during the 150-ms interval post-stimulus appearance. We took this maximum value and subtracted away average baseline activity as our proxy for visual response magnitude. Average response magnitude in SC was 120.3 sp/sec (**Fast**) and 116.7 sp/sec (**Accurate**). We found no evidence for effect of condition on response magnitude (t_(9)_=0.50, p=0.63, BF = 2.9) in SC neurons. Average response magnitude in FEF was 31.7 sp/sec (**Fast**) and 27.9 sp/sec (**Accurate**). Visual response magnitude in FEF tended to be greater in the **Fast** condition, but this effect was not significant (t_(21)_ = 1.67, p = 0.11, BF = 1.4).

Heitz and Schall (2012) did not report visually-responsive activity on error trials. Obviously, in the **Fast** condition more endpoint error trials were collected, and in the **Accurate** condition error trials were less common. For the **Accurate** condition, in many cases error trials were too rare to allow analysis for each neuron. Therefore, we report population averages. In both FEF and SC, endpoint errors occurred when visual salience neurons treated a distractor as if it were the target. We asked whether SAT-related speed constraints affected this process of erroneous distractor selection. For this analysis, in addition to visually-responsive FEF neurons from *Da*, we also analyzed error-related activity in sets of 83 and 90 FEF neurons from monkeys *Q* and *S*.

For each set of neurons, we computed the average normalized SDF for endpoint error trials with response directed either into or outside of the neuron’s movement field (Fig. 8A; *Q*, *S*, and *Da*). We estimated the average saccade-in vs. saccade-out activity for each neuron, and then applied the Mann-Whitney U-test to the set of average SDFs. Average TST for FEF neurons on endpoint error trials in the **Fast** condition was 182 ms (*Q*), 230 ms (*S*), and 226 ms (*Da*); average TST in the **Accurate** condition was 411 ms (*Q*), 437 ms (*S*), and 511 ms (*Da*). Average TST on error trials in SC (Fig. 8B; *Da & Eu*) was 142 ms (**Fast**) and 331 ms (**Accurate**).

**Figure 8.**
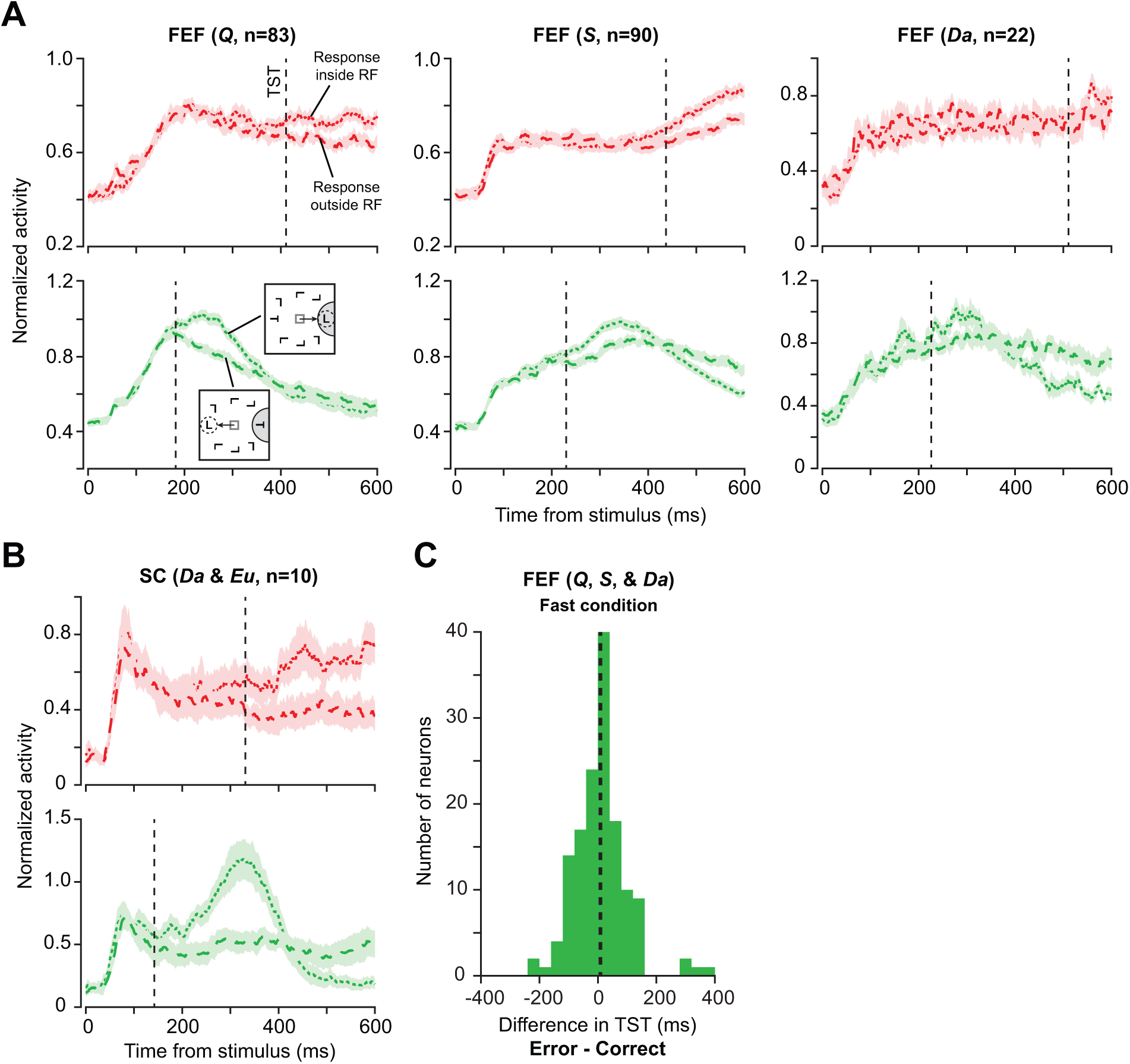
Origin of targeting errors in FEF neural activity. (**A**) Average normalized SDF of all visually-responsive FEF neurons (*Q:* n = 83, *S:* n = 90, and *Da:* n = 22) during the 600-ms interval following presentation of the array stimulus. Only trials with correctly timed and mis-directed responses were included. Density functions for **Accurate** (*top*) and **Fast** (*bottom*) conditions are shown separately. Diagrammatic inset shows schematic of the search array on trials when the erroneous response was directed either into or outside of the neuron’s RF. Black dashed lines show average TST per condition. Average TST in FEF during the **Fast** condition was 182 ms (*Q*), 230 ms (*S*), and 226 ms (*Da*); average TST in the **Accurate** condition was 411 ms (*Q*), 437 ms (*S*), and 511 ms (*Da*). Error regions are SEM. (**B**) Average normalized SDF of all visually-responsive SC neurons (*Da* n = 4, *Eu* n = 6). Average TST in SC was 142 ms in the **Fast** condition, and 331 ms in the **Accurate** condition. Conventions as in (A). (**C**) Distribution of the difference between TST on trials with correct and mis-directed responses. Dashed line shows across-neuron average difference of 8 ms. Distribution includes neurons from monkeys *Q*, *S*, and *Da*. The difference measure could only be computed for neurons that exhibited target selection on both correct and error trials (n = 143).

Was the timecourse of target selection comparable on correct and error trials? To answer this question, for each FEF neuron that selected the target amongst distractors, we estimated TST separately on correct and error trials. We focused this analysis on the **Fast** condition, as endpoint errors in the **Accurate** condition were too sparse to provide a reliable estimate of TST for most neurons. Figure 8C shows the distribution of difference in TST between correct and error trials for all such FEF neurons (n = 143). In FEF, target selection time did not change if the source of selection was an erroneous distractor, instead of the target (paired t-test, t_(142)_=1.1, p=0.28, BF = 5.9). On average, TST was 8 ms later on endpoint error trials.

### Response preparation adjustments

The activity of movement neurons in FEF and SC initiates saccades when a stochastic accumulation of discharge rate reaches a level that is invariant across RT (Boucher et al., 2007; Ding and Gold, 2012; Hanes and Schall, 1996; Pouget et al., 2011; Purcell et al., 2010, 2012b; Ratcliff et al., 2007; Woodman et al., 2008). We found adjustments of the movement neuron accumulation process during SAT.

We first asked whether speed constraints affected buildup activity in FEF. We analyzed 18 FEF neurons with saccade-related buildup activity (*Da*: 16 visuo-movement & 2 pure movement). We normalized buildup activity to the maximum average firing rate during the peri-saccadic time interval from 100 ms before to 100 ms after saccade initiation. Figure 9A shows the average SDF for all 16 visuo-movement FEF neurons. Activity at saccade onset was higher in the **Fast** condition. Figure 9B shows the buildup activity for the 2 pure movement neurons from FEF. Buildup activity of these neurons reached a higher value during trials that emphasized accuracy to speed. Across the population of FEF neurons with saccade buildup activity, the activity at saccade onset was higher in the **Fast** condition (Fig. 9C, paired t-test, t_(17)_ = 2.38, p = 0.03).

**Figure 9.**
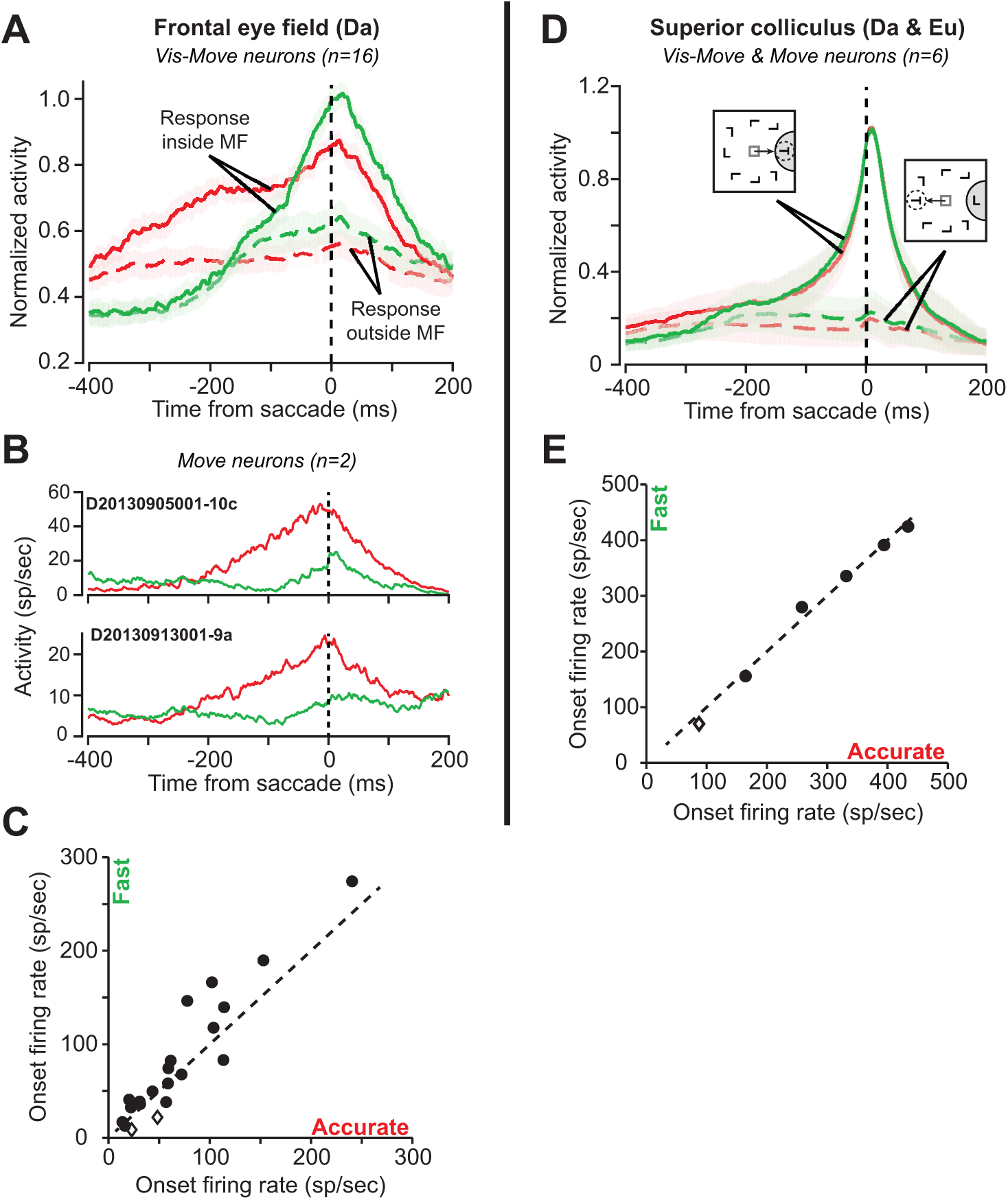
Saccade-related buildup activity of FEF and SC neurons during speed-accuracy tradeoff. (**A**) Average normalized SDF of peri-saccadic buildup activity across all FEF visuo-movement neurons (*Da* n = 16). Solid and dashed lines denote trials with response directed into and outside of the neuron’s MF, respectively. Buildup activity of each neuron was normalized to peak saccade-locked activity (averaged across all trials) during the interval from −100 ms to 100 ms relative to saccade. Activity of visuo-movement FEF neurons at saccade initiation was higher in the **Fast** condition. (**B**) Average SDFs of peri-saccadic buildup activity of the two FEF neurons with pure movement-related activity. Activity of the two neurons at saccade initiation was higher in the **Accurate** condition. (**C**) FEF neuron-wise relationship between activity at saccade onset in the **Fast** and **Accurate** conditions. Each dot represents a single neuron. Dashed line is unity. The two diamonds represent the pure movement neurons. Activity at saccade onset was consistently higher in the **Fast** condition (paired t-test, p = 0.03). (**D**) Average normalized SDF of peri-saccadic buildup activity of all SC neurons with saccade-related activity (*Da* n = 2, *Eu* n = 4). Diagrammatic insets show schematic of the search array on trials with response directed into and outside of the neuron’s MF. Conventions as in (A). (**E**) SC neuron-wise relationship between activity at saccade onset in the **Fast** and **Accurate** conditions. Activity of movement-related SC neurons at saccade initiation was invariant across conditions (paired t-test, p = 0.80).

We next asked whether imposition of speed constraints affected the buildup of saccade-related activity in SC. We analyzed activity from 6 SC neurons with buildup activity (*Da*: 2 visuo-movement; *Eu*: 3 visuo-movement & 1 pure movement). Figure 9D shows the across-neuron average SDF relative to saccade initiation. Unlike FEF, in SC the firing rate at saccade initiation was invariant across conditions (Fig. 9E, paired t-test, t_(5)_ = 0.27, p = 0.80, BF = 2.6).

In summary, across the population of 6 movement-related SC neurons, we found no effect of speed constraints on activity at saccade initiation. However, across the population of 18 movement-related FEF neurons, we found a dissociation of speed-related effects on buildup activity, replicating prior findings (Heitz & Schall, 2012). For visuo-movement neurons (n=16), buildup activity at saccade onset was higher in **Fast** condition; however, for pure movement neurons (n=2), buildup activity was higher in **Accurate** condition.

### Saccadic activity and saccade peak velocity

Peak velocity of saccades varied idiosyncratically across conditions. Additionally, saccade-related buildup activity was invariant across conditions (SC), or sensitive to speed constraints (FEF). We therefore asked whether buildup activity in SC and FEF was related to saccade peak velocity. For each trial, we computed the correlation between saccade peak velocity and the average value of the single-trial SDF during the 30-ms interval post-saccade initiation. We chose this interval because most task-relevant saccades had a duration of ~ 30 ms. Across 6 SC neurons with saccade-related activity, we observed a positive correlation between peak velocity and average firing rate for 4 neurons (Pearson correlation coefficient: **Fast** condition 4/6 cells, p<0.05; **Accurate** condition 2/6 cells, p<0.05). Figure 10 shows this relationship for 1 of the 2 neurons with a significant correlation in both conditions (**Accurate**: R = 0.39, p = 9×10^−10^; **Fast**: R = 0.27, p = 0.001). Across 18 FEF neurons with saccade-related buildup activity, we observed a positive correlation for 4 neurons (**Fast** condition: 4/18 cells, p<0.05; **Accurate** condition: 1/18 cells, p<0.05, Pearson correlation coefficient). In summary, we did not observe a systematic correlation between peak velocity and firing rate during saccade, either in SC or FEF. However, we did observe correlations at the single-neuron level in both SC (4/6) and FEF (4/18).

**Figure 10.**
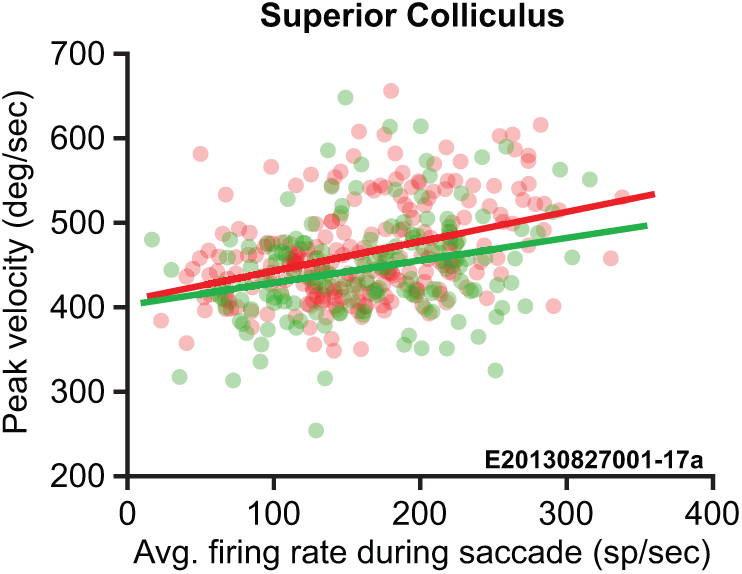
Peak velocity and saccade activity in SC. Relationship between saccadic peak velocity and maximum firing rate for representative SC neuron from monkey *Eu*. Each point represents a single trial. Red and green dots represent data from **Accurate** and **Fast** trials, respectively. Peak velocity of saccades increased with average saccade-related activity for this neuron (Pearson correlation coefficient: **Accurate** R = 0.39; **Fast** R = 0.27). We observed a positive correlation between maximum firing rate and saccadic peak velocity during at least one condition for 4 out of 6 SC neurons with saccade-related activity.

## Discussion

Although this paper reports the fifth neurophysiological study of SAT (Hanks et al., 2014; Heitz and Schall, 2012; Thura and Cisek, 2016, 2017), it is a first in several respects. It is the first replication of an SAT task with new monkeys, demonstrating the reliability of this experimental approach in both behavioral and neural terms. We show that the major performance measures are replicated across four monkeys. These performance measures include the first description of the relationship between response time and the timing deadlines used to enforce the SAT conditions. Also, we report how error rate varies systematically with response time within SAT conditions. These relationships are necessary information for future models of SAT in this task.

This paper is the first detailed description of the variation of saccade dynamics with SAT conditions, monkeys and response time. The idiosyncratic and systematic variation of saccade dynamics contradicts a basic assumption motivating an integrated accumulator explanation of FEF movement neuron modulation during SAT (Heitz & Schall 2012). Thus, more elaborate models will be needed to understand how saccade-related discharges in FEF and SC relate to saccade dynamics.

This paper is the first description of neural adjustments in SC with visual search SAT. We believe these results merit publication because the unique data address an important general question. Confidence in the reliability of the conclusions can be high for the following reasons. First, the performance of the two additional monkeys replicated that of the original two monkeys. Second, the sample of visual salience neurons in FEF of a third monkey replicated major observations reported previously, and the observation of movement neurons with activation at RT higher in **Accurate** relative to **Fast** trials is theoretically very important. Third, the general pattern modulation of SC neurons during visual search replicates previous observations by multiple laboratories (e.g., McPeek and Keller, 2002; Shen and Paré, 2007; White and Munoz, 2011; White et al., 2017). Fourth, each new observation was replicated across monkeys and neurons. Thus, while a larger sample of neurons would increase statistical power, given the well-known, consistent patterns of SC modulation, it is unlikely that the results would change. These new data confirm earlier reports that at the neurophysiological level, SAT is accomplished by a multitude of mechanistically distinct adjustments. As in FEF, when accuracy was cued, proactive baseline discharge rate in SC was reduced. We demonstrate that this modulation occurs immediately upon SAT cue changes and becomes more efficient in successive trials. Unlike FEF, visual responses in SC to the search array did not vary with SAT condition; however, like FEF, the target selection process took more time in the **Accurate** compared to the **Fast** condition. Also unlike FEF, the presaccadic movement activity at saccade onset was invariant across task conditions. Of note, two movement neurons were recorded in FEF of the new monkeys, and both neurons exhibited higher saccade-related activity on **Accurate** as compared to **Fast** trials. The reason for these differences across monkeys and, possibly, structures will require more investigation.

Finally, this paper offers the first description of the origin of errors in SAT through incorrect representation of the evidence. As observed previously (Heitz et al., 2010; Thompson et al., 2005), search errors occurred when visual neurons in FEF and SC treated a distractor as if it were the target.

The general theoretical implications of these SAT neurophysiology data have been detailed before (Heitz and Schall, 2012, 2013). The specific implications of these new observations will be worked out through formal computational models of how salience evidence can be accumulated to guide gaze (Purcell et al., 2010, 2012a).

## Acknowledgments

This work was supported by F32-EY019851 to R.P.H., T32-EY007135 for T.R.R., and R01-EY08890, P30-EY08126, U54-HD083211 and by Robin and Richard Patton through the E. Bronson Ingram Chair in Neuroscience. We thank J. Easley, M. Feurtado, M. Maddox, S. Motorny, J. Parker, M. Schall, and L. Toy for animal care and other technical assistance. Requests for materials should be addressed to JDS.

## Conflict of Interest

The authors declare no competing financial interests.

